# A computational model to understand mouse iron physiology and diseases

**DOI:** 10.1101/323899

**Authors:** Jignesh H. Parmar, Pedro Mendes

## Abstract

It is well known that iron is an essential element for life but is toxic when in excess or in certain forms. Accordingly there are many diseases that result directly from either lack or excess of iron. Iron has also been associated with a wide range of other diseases and may have an important role in aging. Yet many molecular and physiological aspects of iron regulation have only been discovered recently and others are still awaiting elucidation. In the last 18 years, after the discovery of the hormone hepcidin, many details of iron regulation have become better understood and a clearer picture is starting to emerge, at least in qualitative terms. However there is still no good quantitative and dynamic description of iron absorption, distribution, storage and mobilization that agrees with the wide array of phenotypes presented in several iron-related diseases. The present work addresses this issue by developing a mathematical model of iron distribution in mice that was calibrated with existing ferrokinetic data and subsequently validated against data from a series of iron disorders, such as hemochromatosis, β-thalassemia, atransferrinemia and anemia of inflammation. To adequately fit the ferrokinetic data required including the following mechanisms: a) the role of transferrin in deliving iron to tissues, b) the induction of hepcidin by high levels of transferrin-bound iron, c) the ferroportin-dependent hepcidin-regulated iron export from tissues, d) the erythropoietin regulation of erythropoiesis, and e) direct NTBI uptake by the liver. The utility of such a model to simulate disease interventions was demonstrated by using it to investigate the outcome of different schedules of transferrin treatment in β-thalassemia. The present model is a successful step towards a comprehensive mathematical model of iron physiology incorporating cellular and organ level details.

## Introduction

Iron is an essential metal in living organisms, which is required as a co-factor for many proteins with roles in electron transport and oxygen binding. In mammals the majority of the body’s iron is used for oxygen transport in the form of hemoglobin in red blood cells (RBC). Anemia arises whenever there is an impairment of functional hemoglobin, as is the case with low iron availability for RBC production, and results in debilitating low energy due to decreased oxygen delivery. Anemia can be caused by iron deficient diets or from disregulation of the body’s iron distribution. Iron can also be extremely toxic if present in labile forms (*i.e*. free or weakly bound) as it can catalyze synthesis of hydroxyl radical, one of the most potent oxidants known, possibly the cause of iron being associated with a large number of apparently unrelated diseases, such as cardiovascular, neurodegenerative, cancer, and many others (Kell, 2009). Thus mechanisms have evolved to control the total iron content in the body, its distribution across organs, and to limit its presence in labile forms. Some of these mechanisms have been known for decades, but some important aspects are only now being unraveled (e.g. Kautz *et al*, 2014).

The present understanding of mammalian iron regulation reveals an intricate network containing many feedback loops (Hower *et al*, 2009; Wallace, 2016). Dietary iron is absorbed by duodenal enterocytes which release it to the blood where it binds to transferrin (Tf), a transport protein with a very strong affinity for ferric iron at neutral pH. Transferrin-bound iron is delivered to cells through binding of this protein to specific receptors with subsequent internalization, acidification-driven iron release, and recycling of the apo-transferrin back to blood. In normal conditions the amount of plasma non-transferrin bound iron (NTBI) is extremely low. This is important because NTBI is able to catalyze the Fenton reaction and produce hydroxyl radical.

Iron is required by all tissues for its cofactor role in several enzymes, though quantitatively this is a very small proportion of the total iron in adult animals. The majority of absorbed iron is used for hemoglobin and circulates between erythroid precursor cells in the bone marrow, RBC and macrophages in the spleen. When there is iron overload, either due to excessive dietary intake or as a consequence of disease, the excess iron accumulates predominantly in the liver.

Systemic iron regulation happens primarily through the control of iron export from cells, a process mediated solely by ferroportin (Ganz, 2005) and regulated by the peptide hormone hepcidin (Ganz, 2013). Hepcidin inhibits ferroportin activity resulting in iron immobilization inside cells, and this also inhibits iron acquisition from the diet because it impedes enterocytes from delivering newly acquired iron to the plasma. Hepcidin secretion by the liver is transcriptionally stimulated by high plasma transferrin-bound iron and, indepedently, also by interleukin-6. The latter mediates an important defense mechanism against microbial growth by reducing the circulating iron, trapping it in organs through maintenance of high levels of hepcidin. As the plasma iron increases, so does the concentration of hepcidin, and this forms a negative feedback loop on dietary iron acquisition. Mutations that impair sensing of plasma iron by the liver or the activity of hepcidin result in hemochromatosis, a disease characterized by very high levels of total body iron, with a strong accumulation in the liver. Untreated, hemochromatosis can lead to liver disease, cancer, and cardiovascular disease (Franchini & Veneri, 2005). Chronic inflammation upregulates hepcidin production resulting in a type of anemia where, even though the total body iron is not low, its availability for production of new RBC is impaired (Ganz, 2003). In this type of anemia of inflammation, iron is trapped in the liver and spleen, and absorption of dietary iron is inhibited, resulting in inadequate iron supply to the bone marrow and consequently low levels of hemoglobin.

Since one of the largest uses of iron in the body is in the production of hemoglobin it is relevant to consider the regulation of erythropoiesis. The hormone erythropoietin (EPO), secreted by the kidneys, stimulates the proliferation of erythroid progenitor cells and therefore increases the consumption of iron by the bone marrow, and consequently its level in RBC. EPO is increased under hypoxia and abrogated when oxygen is plentiful. A recently discovered hormone, erythroferrone (Kautz *et al*, 2014), couples EPO to hepcidin signaling. Erythroferrone secretion by erythroid precursors is stimulated by elevated EPO levels, and erythroferrone inhibits the secretion of hepcidin – resulting in a negative regulation of hepcidin by EPO. EPO is elevated in conditions that require increased production of RBC, such as after hemorrhage or in anemia, resulting in increased erythroferrone and finally in suppression of hepcidin (Kautz *et al*, 2015; Kim & Nemeth, 2015; Jiang *et al*, 2016), allowing iron to be mobilized from the liver and spleen to the plasma, and also increasing dietary iron absorption. This provides the required increased levels of iron for increased RBC production.

Complex biological phenomena are difficult to understand due to the large number of molecular interactions that often have counter-intuitive outcomes. Their understanding requires a systems-level and quantitative analysis such as aided by computational models (Kitano, 2002). Computational models are therefore useful tools for understanding the mechanisms contributing to homeostasis and disease (Glynn *et al*, 2014; Brodland, 2015). Iron regulation, as briefly described above, is sufficiently complex to warrant such an analysis. Several models have been previously developed that focus on various aspects of mouse iron homeostasis at the whole-body level (Lopes *et al*, 2010; Enculescu *et al*, 2017; Parmar *et al*, 2017). However, these models are not capable of adequately explaining the iron distribution in conditions of iron overload with the same set of parameters as for iron deficiency. A recent model focused on chronic kidney disease (Sarkar *et al*, 2018), while useful for the purpose intended, is not applicable to many other iron-related diseases (and does not include the processes of iron absorption and excretion). Our previous work revealed two major problems with model predictions in high iron conditions: a) an overestimation of iron levels in the RBC, and b) did not predict iron accumulation in the liver (Parmar *et al*, 2017). However, data shows that under dietary iron overload mice do accumulate the excess iron in the liver, not in the RBC. This suggested to us that the regulation by hepcidin alone is not sufficient and that other mechanisms play a role in regulating iron distribution in excess iron conditions. The present work presents an improved model of systemic mouse iron homeostasis that is capable of explaining iron distribution in wide ranges of total body iron. The results communicated here suggest that the regulation of iron distribution depends on the combined action of hepcidin and erythropoietin (and implicitly also erythroferrone), with an additional important role of the direct import of NTBI into the liver.

Since models can be deceiving, particularly when used for extrapolations, it is important to assess their validity by applying them to a domain different from that used to construct them (Carusi, 2014; Patterson & Whelan, 2017). We carried out validation of this new model by using it to successfully predict the phenotype of a number of iron disorders, several of which result in iron accumulation. This is a clear improvement over previous systemic iron distribution models. The results show that the model can be used to aid our understanding of the pathophysiology of various iron disorders and different dietary regimes. The model was also applied to predict the outcome of a proposed treatment of beta-thalassemia with transferrin injections, and the difference between intravenous and intragastric iron supplementation.

## Methods

All simulations and computational analyses were carried out with the open source software COPASI versions 4.20 to 4.23 (Hoops *et al*, 2006; Bergmann *et al*, 2017). Model calibration was carried out by non-linear least-squares as implemented in COPASI and detailed below. Model validation was carried out for a series of physiological and disease conditions different from those used for parameter estimation (see Results section). Validation is important to establish generality of the model and as a check that the parameter estimation did not significantly overfit the data.

### Model

The model was constructed based on a previous version (Parmar *et al*, 2017) with important modifications as described below. Full details of the model are included in the supplementary information: equations, parameter values (Table S1), and initial conditions (Table S2). A diagram using the SBGN standard (Le Novère *et al*, 2009) and constructed with the aid of the software CellDesigner (Funahashi *et al*, 2003) is depicted in Figure S1. Simulation files in the COPASI format and in the SBML standard (Hucka *et al*, 2003) are also included in the supplementary information. Two versions of the model (with and without radioactive iron tracer species) have been submitted to the BioModels database (Chelliah *et al*, 2015) where they are available with identifiers MODEL1805140002 and MODEL1805140003.

Following our previous work, we estimate parameters based on radioactive iron tracer data with a version of the model where each reaction or transport step can happen with radioactive or nonradioactive iron species; all kinetic constants are assumed to be the same irrespective of iron isotope. The model parameters were adjusted to match simultaneously data from three tracer experiments with mice subjected to high, low, and adequate iron diets. Since the tracer data are only important for estimation of parameter values, the model version with radioactive species was only used for parameter estimation; model validation and other results were obtained with the version without tracer, except where noted. The model considers seven compartments with fixed volumes (determined as in Parmar *et al*, 2017), namely: duodenum, plasma, liver, bone marrow, red blood cells (RBC), spleen, and rest of body (accounting for the remaining parts of a mouse body). Below we discuss the specific new features that were crucial to making the present model pass a wide array of validation data.

#### NTBI uptake by the liver

It is widely accepted that the main form of iron import into liver cells, as with most other cell types, is through transferrin and the transferrin receptors. Plasma non-transferrin bound iron (NTBI) is normally at such low concentrations that it is hard to quantify, but under high iron loading this species can increase considerably due to transferrin saturation and the excess iron ends up mostly in the liver. The previous version of our model (Parmar *et al*, 2017) was unable to accumulate iron in the liver under a high iron diet and we noted that under that condition the model showed an unreasonably large accumulation of NTBI. Independent fits of that model against the high iron and adequate iron diets differed more in the liver iron import than any other parameters. This led us to the hypothesis that liver uptake of NTBI must be important. Indeed there is experimental support for this hypothesis in experiments showing the membrane protein ZIP14 (SLC39A14) is able to uptake iron into cells (Liuzzi *et al*, 2006) and its repression results in reduced liver iron uptake in hemochromatosis (Liuzzi *et al*, 2006; Jenkitkasemwong *et al*, 2015). Nam and colleagues observed that liver ZIP14 expression is elevated under high iron loads (Nam *et al*, 2013). Therefore we added a new transport step in the model allowing NTBI to be directly taken up by the liver. In order to allow this transport rate to increase in high iron conditions more steeply than a simple Michaelis-Menten rate law, we used instead a substrate activation kinetic rate law:

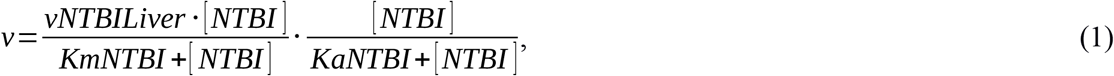

where *vNTBILiver* is the limiting rate (Vmax), *KmNTBI* is the Michaelis constant, and *KaNTBI* is an activation constant. In the case of the model with a radioiron tracer, we need to consider the two NTBI species (normal and radioactive); the two work together as activators, but compete as substrates, thus the rate law becomes:

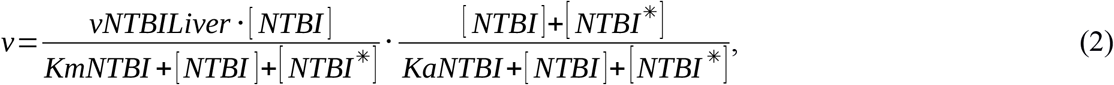

for the liver NTBI uptake reaction, and

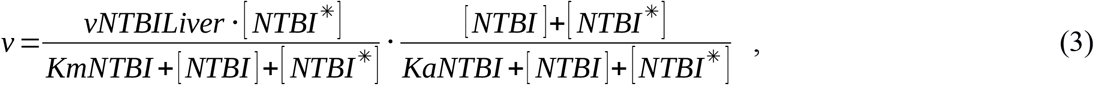

for the liver uptake of NTBI* (the corresponding radioactive tracer species).

#### Erythropoetin

Because our previous model overestimated the amount of RBC iron in a high iron diet, we modified the model under the hypothesis that the regulation of erythropoiesis is important for iron distribution. The hormone erythropoietin (EPO), secreted in the kidneys, stimulates the proliferation of erythroid progenitor cells in the bone marrow. EPO secretion is inversely proportional to oxygen availability in the RBC, which is expected to be proportional to the amount of RBC iron in the model (*FeRBC*). Thus, as *FeRBC* increases, EPO decreases and erythropoiesis slows down.

This process of EPO control of erythropoiesis was included in the model by addition of a new species, *EPO* (located in the plasma), which has a synthesis rate inhibited by *FeRBC* according to a Hill function:

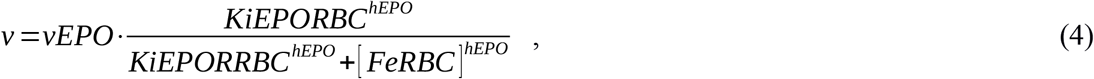

where *vEPO* is a constitutive synthesis rate, *KiEPORBC* is an inhibition constant for the RBC iron, and *hEPO* a Hill coefficient. The rate of degradation of EPO is represented by a simple first order process.

The effect of EPO on erythropoiesis was then added by having *EPO* stimulate *a)* the transport of transferrin-bound iron into the bone marrow (because more erythroid precursor cells are importing iron), and *b)* the transfer of iron from bone marrow to RBC (because erythroid precursors are differentiating into mature RBC at a higher rate). The stimulating effect of *EPO* on these two transport steps was made linear, according to the rate law:

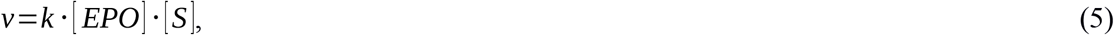

where *k* is a constant, and *S* is the substrate (a transferrin-bound iron species for the reactions of iron import to *FeBM*, or the transfer of iron from bone marrow to RBC); *k*‧[*EPO*] has units of a first order rate constant. (*k* becomes *kInBM* for the transfer of iron from transferrin to bone marrow, or *kInRBC* for the transfer of iron from bone marrow to RBC.)

The regulation conferred by EPO thus provides a negative feedback loop from *FeRBC* to its own synthesis, stabilizing this variable against changes of total body iron.

#### Hepcidin

The hepcidin synthesis rate depends on the level of transferrin-bound iron and is also inhibited by EPO. Thus we used the following equation for the rate of hepcidin synthesis:

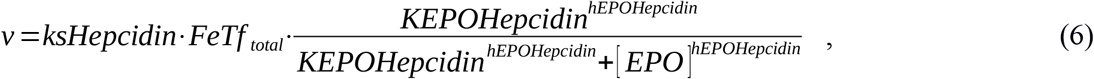

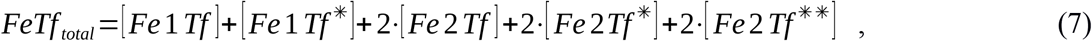

where, *ksHepcidin* is a linear activation constant, *FeTf*_*total*_ is the total concentration of iron bound to transferrin molecules, *KEPOHepcidin* is an inhibition constant, and *hEPOHepcidin* is a Hill coefficient. The first term in this equation represents the sensing of transferrin-bound iron, while the second term represents the inhibitory effect exerted by EPO on hepcidin synthesis. The latter effect has been shown to be mediated by erythroferrone (Kautz *et al*, 2014, 2015; Jiang *et al*, 2016), though the present model only includes the overall effect without representing erythroferrone explicitly.

### Simulations and parameter estimation

To estimate the values of the parameters, the model was set up to simulate the same protocol taken in the ferrokinetic experiments that generated the calibration data (Schümann *et al*, 2007; Lopes *et al*, 2010). Briefly these consisted of three mice groups which were put on different iron diets (iron-deficient, adequate, and iron-rich) for 5 weeks, then injected with a tracer and followed for a 28 day time course, with the radioactive tracer quantified in all their organs. To replicate this protocol the simulation happened in two phases: 1) phase one starts by setting the value of the *vDiet* parameter to simulate the three iron diets; this phase lasts 5 weeks as in the experiments. 2) The second phase starts with the addition of a bolus of radioactive iron (setting the total radioactivity in the variable *NTBI*\* with a discrete event) and is simulated for another 28 days. It is only in phase two that the tracer data is compared between simulation and experiment and their distance minimized using least squares. The regression was carried out by minimization of the sum of squares of the residuals using COPASI’s parameter estimation task. This was effected first by application of a global optimizer (SRES, Runarsson & Yao, Xin, 2000) followed by a local optimizer (Hooke & Jeeves, 1961).

The values of kinetic constants obtained from this procedure are those that best fit the tracer time courses for all three mice groups *simultaneously* (adequate, deficient, and rich diets). To ensure that the initial conditions of the model correspond to a steady state (as the mice are expected to have been in a quasi steady state), the model includes specific constraints consisting of algebraic expressions relating parameter values and the initial concentrations that were derived from the right-hand side of the set of ODEs.

### Validation

The calibrated model is then put to test against a variety of experiments carried out with mutant or diseased mice previously reported in the literature. These simulations are carried out with the calibrated parameter set, and no more fitting is carried out. Some parameter values are changed corresponding to the specific mutations or their known physiological outcomes, and these are indicated in the text in each specific case. Therefore the model is being tested against independent experiments that form a robust validation procedure demonstrating the wide applicability of the model (Carusi, 2014) and a low degree of overfitting. The simulation protocols are specific for each case and are described together with the results.

## Results

A single parameter set was obtained that is able to fit simultaneously the three data sets with different iron diet (Table S1). The only parameter value that is different for each data set is the rate of entry of iron into the duodenum (*fDiet‧vDiet*), a consequence of the different iron content in the diets. Figure 1 compares the model fit with the observations for the liver and RBC (Figures S2-S4 depict data for all the compartments). The model reaches a stable steady. In the case of the adequate iron diet, the steady state values of total iron, Tf saturation, hepcidin and EPO for the adequate diet (Table S2) are in the range observed in the literature (Ehrenstein & Hevesy, 1959; Ajioka *et al*, 2002; Murphy *et al*, 2007; Xing *et al*, 2008; Morán-Jiménez *et al*, 2010). As expected, in the iron-deficient diet the Tf saturation, total iron, and hepcidin concentration are lower than in the standard iron diet, while opposite changes were observed in the iron-rich diet.

**Figure 1.**
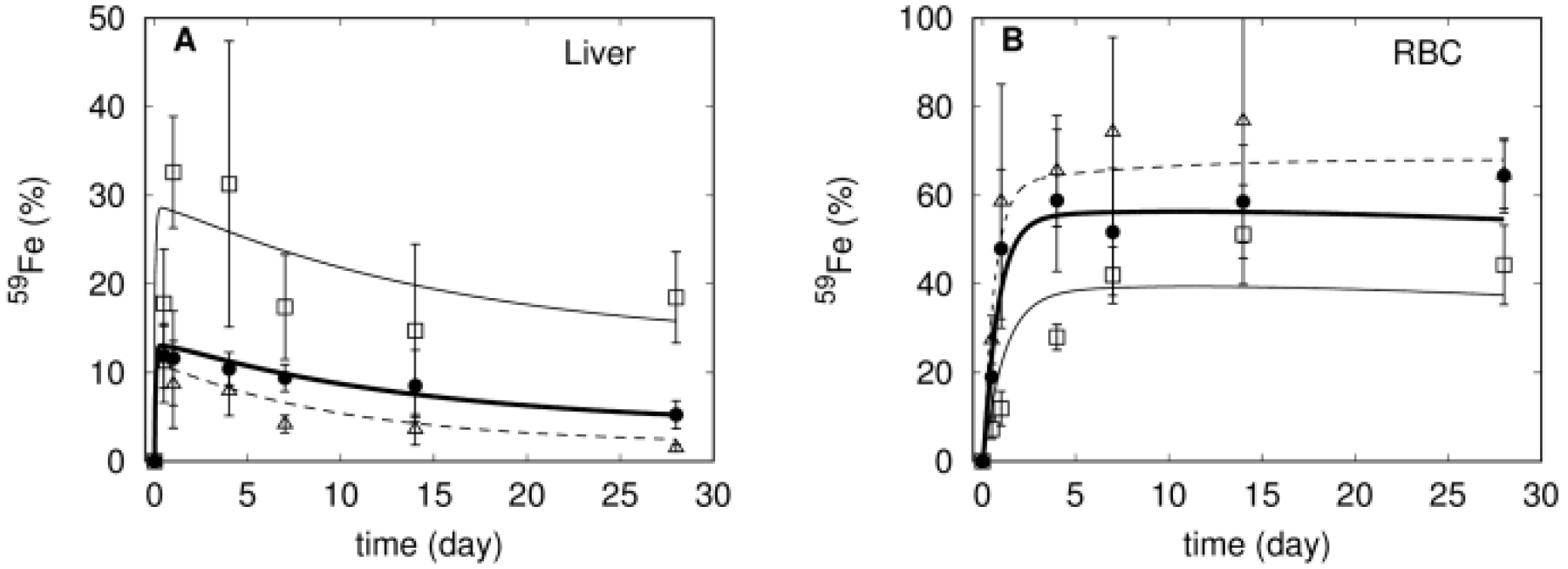
Comparison of ^59^Fe level and model fit for the liver (A) and RBC (B) compartments. Filled circles and bold line for the experimental data and model fit of adequate diet, squares and thin line for experimental data and model fit of iron rich diet, and triangles and dashed line for experimental data and model fit of iron deficient diet.

One of the main goals of a model is to better understand the experimental observations. This is the case here, particularly as the model includes predictions of the levels of the non-radioactive iron species and hormones, which were not measured experimentally. Two important features are revealed by observation of the behavior of the model in the 5 weeks prior to the tracer injection. In the iron-rich diet, excess iron accumulates in the liver and this is explained by considering the direct transport of NTBI into this compartment through the ZIP14 transporter. We observe that during the first five weeks under this diet transferrin becomes very close to saturation (Fig. 2A) resulting in a build up of NTBI and liver iron (Fig. 2B). An unexpected observation in the experimental data set is that under the iron-rich diet the radioactive tracer reaches lower levels in the RBC than in the other two dietary regimes (Fig. 1B). But the model is able to provide an explanation of why this apparently counter-intuitive result happens. Inspecting the simulated time course of the 5 weeks of the iron-rich diet prior to the tracer injection reveals that there is increased non-radioactive iron in the RBC ([*FeRBC*]) and this in turn reduces [*EPO*] (Fig. 2C). When the ^59^Fe is injected, its entry to the bone marrow and transfer to RBC is now restricted by the lower [*EPO*], thus resulting in a lower percentage of ^59^Fe accumulation in RBC than in the adequate and deficient diets. This result confirms the need to include EPO in the model, as otherwise the simulation of the tracer dynamics in the RBC would not match the observations.

**Figure 2.**
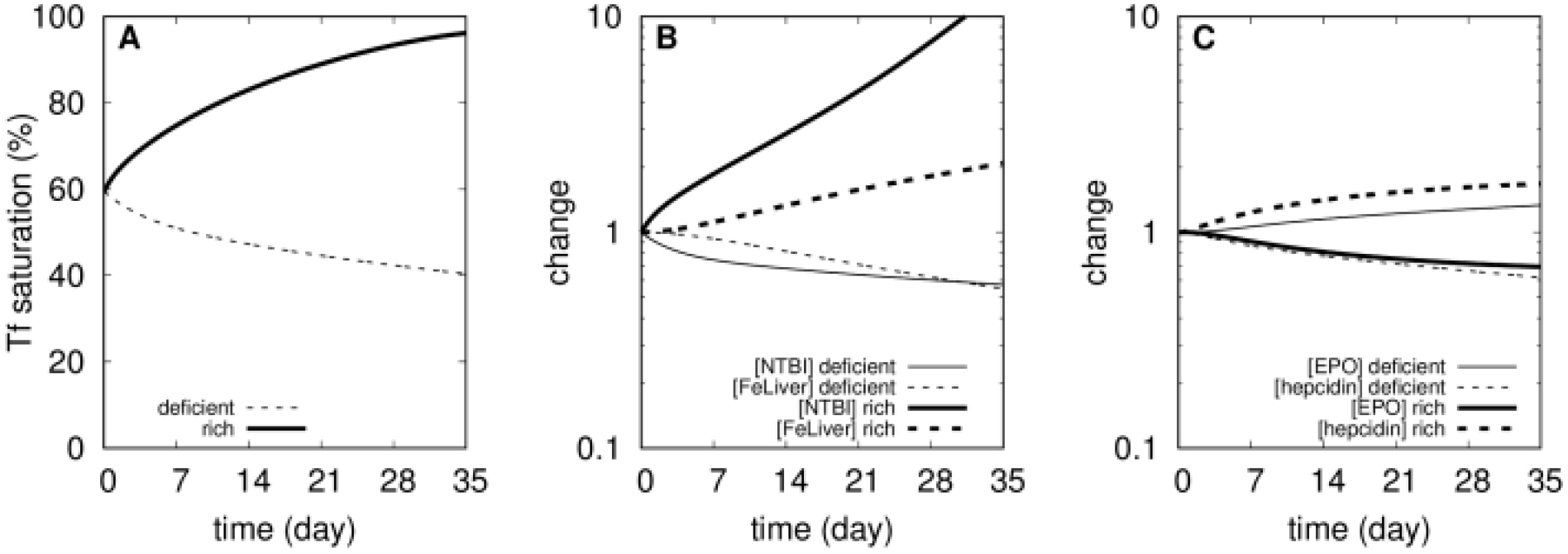
Simulated physiological changes during the first 5 weeks under deficient and rich iron diets. A – Changes in transferrin saturation. B – Changes in [*NTBI*] and [*FeLiver*], relative to their value in the adequate diet (time zero). C – Changes in [*EPO*] and [*hepcidin*], relative to their value in the adequate diet (time zero).

With this successful fit to the ferrokinetic data under three iron dietary regimes, we then apply the resulting model to simulate various pathological conditions that result in iron disregulation, and show the extent to which it is able to predict the corresponding phenotypes, and therefore validated.

### Hereditary hemochromatosis

Hereditary hemochromatosis (HH) is a very common genetic defect among Caucasians that leads to iron overload. HH is caused by genetic disruption of the hepcidin-ferroportin regulatory mechanism. HH can originate from mutations in genes involved in the signaling pathway that induces hepcidin expression as a function of transferrin-bound iron, the most common being HFE (Feder *et al*, 1996), but also TFR2 (Camaschella *et al*, 2000) and HJV (Papanikolaou *et al*, 2004). Mutations in the hepcidin gene (HAMP) cause a rare type of juvenile hemochromatosis (Roetto *et al*, 2003). All of these mutations lead to low levels of hepcidin activity and therefore higher ferroportin activity, resulting in higher absorption of dietary iron through the duodenal enterocytes. The predominant phenotypic alteration in HH is high level of total body iron with accumulation in various tissues, but predominantly in the liver. The severity of iron overload increases in the order from HFE^-/-^ to TFR2^-/-^ to HJV^-/-^ or HAMP^-/-^. Here we use the computational model to simulate hemochromatosis; this is achieved by setting the rate of synthesis of hepcidin (*ksHepcidin*) to zero. This is similar to an induced knockout of the HAMP gene in a normal mouse (*e.g*. by action of a siRNA, or a drug that would block hepcidin activity). We compare the mutant phenotype with the control after 365 days (Table 1). With the HAMP mutation the model does not achieve a steady state and accumulated unlimited amount of iron in the liver compartment (and consequently the total body iron), reflecting the large liver iron accumulation observed in untreated HH (Zhou *et al*, 1998; Subramaniam *et al*, 2012; Jenkitkasemwong *et al*, 2015). In the model transferrin is essentially fully saturated, which is in agreement with observations in animal models of this disease (Zhou *et al*, 1998; Subramaniam *et al*, 2012; Jenkitkasemwong *et al*, 2015), and the level of NTBI is significantly elevated (Jenkitkasemwong *et al*, 2015). Spleen iron is lower than normal, also seen in animal models (Subramaniam *et al*, 2012; Jenkitkasemwong *et al*, 2015). No major change was observed in the RBC iron content despite the high plasma iron level, which is also in agreement with experimental observations (Zhou *et al*, 1998). In summary the model reflects all of the experimental observations of mice models of hemochromatosis.

**Table 1.** Hemochromatosis simulation. The computational model was modified to suppress hepcidin synthesis (HAMP^-/-^) and variables were quantified after 365 days. Change is measured as the ratio HAMP^-/-^/WT. Experimental support refers to publications where similar changes have been observed experimentally in mouse models.

| Variable | WT | HAMP<sup>-/-</sup> | Change | Experimental support |
| --- | --- | --- | --- | --- |
| [FeDuo] (mM) | 10.0 | 2.04 | 0.20 × |  |
| [FeRBC] (mM) | 23.0 | 24.5 | 1.1 × | (Zhou *et al*, 1998) |
| [FeBM] (mM) | 2.08 | 3.35 | 1.6 × |  |
| [FeSpleen] (mM) | 2.27 | 0.532 | 0.23 × | (Subramaniam *et al*, 2012; Jenkitkasemwong *et al*, 2015) |
| [FeLiver] (mM) | 1.30 | 20.8 | 16 × | (Zhu *et al*, 2008; Subramaniam *et al*, 2012; Jenkitkasemwong *et al*, 2015) |
| [NTBI] (nM) | 40.0 | 8,839 | 221 × | (Jenkitkasemwong *et al*, 2015) |
| [FeRest] (mM) | 0.402 | 0.228 | 0.57 × |  |
| [hepcidin] (nM) | 23.0 | 0 |  |  |
| [EPO] (pM) | 2.43 | 1.60 | 0.66 × |  |
| TF saturation (%) | 59.5 | 99.8 | 1.7 × | (Zhu *et al*, 2008; Subramaniam *et al*, 2012; Jenkitkasemwong *et al*, 2015) |
| Total Fe (mg) | 1.60 | 2.74 | 1.7 × |  |

### Effect of transferrin saturation on plasma iron clearance

Craven *et al.* (1987) observed that the clearance of intravenously given ^59^Fe from plasma was much faster in animals that had their transferrin previously saturated with iron compared with controls with a normal level of transferrin saturation. Additionally, ^59^Fe accumulated mainly in the liver and pancreas when transferrin was fully saturated, otherwise it appeared primarily in the bone marrow, spleen, and RBC (Craven *et al*, 1987; Jenkitkasemwong *et al*, 2015). These experiments were simulated with the model by first bringing the transferrin saturation to near 100% through the addition of an appropriate amount of NTBI. This was then followed by an addition of radioactive tracer in the form *NTBI*\*. The simulated tracer time course was then compared to a control simulation where the radioactive tracer is injected without first saturating transferrin. The simulation results (Fig. 3A-B) show a faster clearance of the plasma ^59^Fe when transferrin is pre-saturated than without it, in line with the experimental results of Craven *et al.* (1987, see their Fig. 3). The simulation also shows that the liver acquires a larger proportion of the tracer when transferrin is pre-saturated than in the control case, which is also in line with the experimental data (Craven *et al*, 1987).

**Figure 3.**
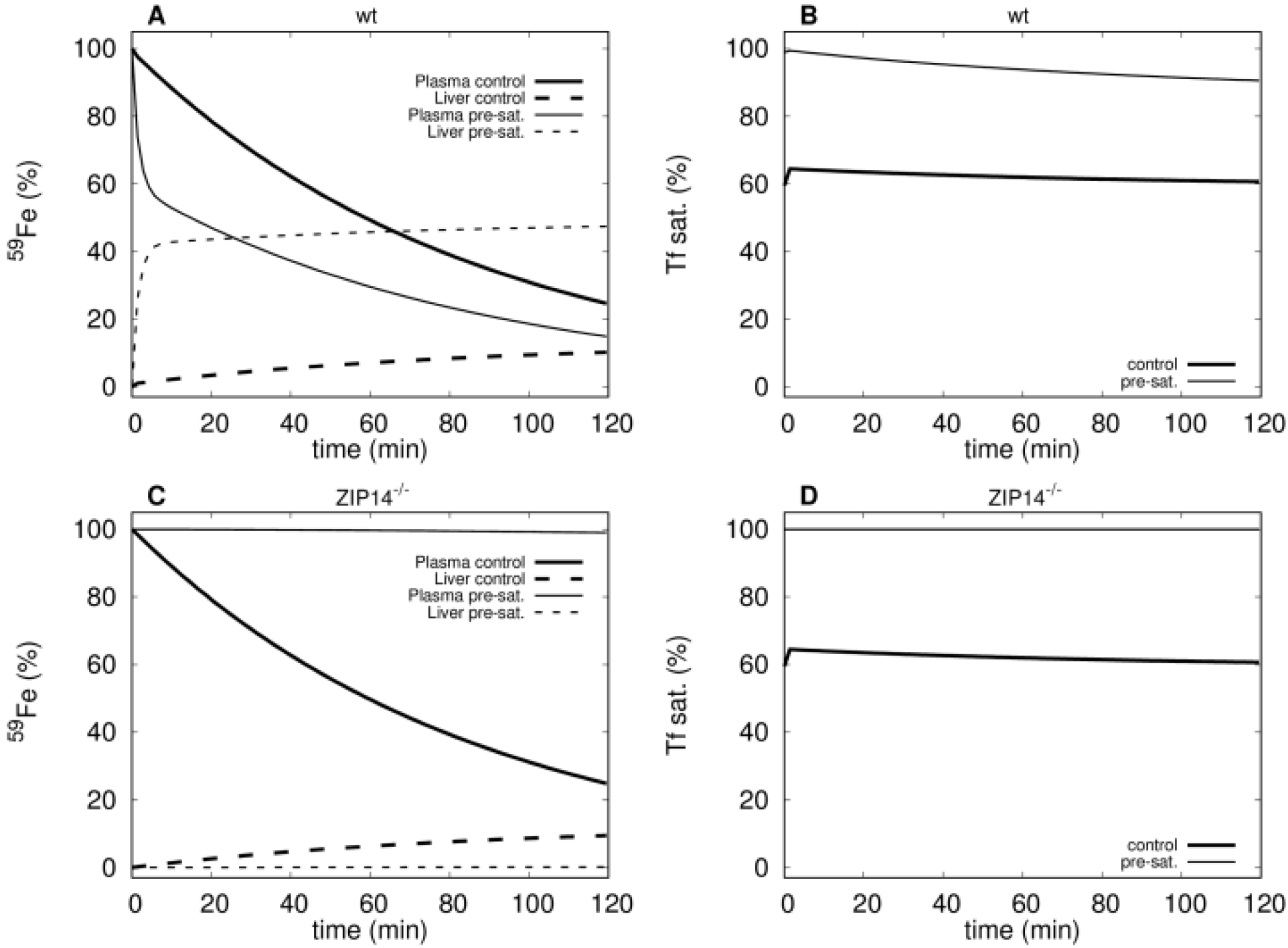
Clearance of ^59^Fe tracer with and without prior saturation of transferrin in wild type and Zip14^-/-^ mutants. **A** – Tracer levels in plasma and liver in the control (bold lines) and immediately after a full saturation of transferrin (thin lines) in the wild type mouse; note the faster clearance from plasma when Tf is pre-saturated and simultaneously higher accumulation in the liver. **B** – Transferrin saturation in control (bold line) and in the pre-saturated treatment (thin line) in the wild type mouse. **C** – Tracer levels in plasma and liver in the control (bold lines) and immediately after a full saturation of transferrin (thin lines) in the Zip14^-/-^ mouse; note the persistence of tracer when Tf is pre-saturated and no accumulation in the liver. **D** – Transferrin saturation in control (bold line) and in the pre-saturated treatment (thin line) in the Zip14^-/-^ mouse.

### Contribution of NTBI uptake by the liver

We now investigate how the model behaves if the direct NTBI uptake by the liver is removed. Thus we simulate the experiments carried out by Jenkitkasemwong *et al.* (2015) where they knocked out the gene for the ZIP14 transporter. They observed that the liver accumulation of ^59^Fe under conditions of saturated transferrin could be prevented by disruption of the ZIP14 transporter. They observed that the kidney and spleen acquired more iron in ZIP14^-/-^ mice compared to controls when transferrin was saturated. This behavior can indeed be reproduced with the present model by setting the rate parameter for liver uptake of NTBI to zero (simulating ZIP14^-/-^). Figure 3C shows that liver iron accumulation is essentially eliminated, and indeed we also observed a higher iron accumulation in the spleen and “rest of body” compartments. There was no effect of removing NTBI uptake on the transferrin-bound ^59^Fe distribution in agreement with the experiments.

Jenkitkasemwong *et al.* also investigated the effect of the ZIP14 knockout in mouse models of HH (HFE^-/-^:ZIP14^-/-^ and HJV^-/-^:ZIP14^-/-^) which greatly reduced the accumulation of iron in the liver (Jenkitkasemwong *et al*, 2015). To reproduce this result with the model, we set the rate of liver uptake of NTBI to zero (ZIP^-/-^) while simultaneously setting the rate of hepcidin synthesis to zero (HAMP^-/-^, causing HH). The results in Table 2 show that this prevented iron accumulation in the liver, consistent with the experimental observations (Jenkitkasemwong *et al*, 2015). However, the model did not reproduce the significant iron accumulation in the spleen and the “rest of body” (which is where the model includes kidney, heart and other organs) as was observed in the experiments. Instead the model showed a very large increase of NTBI. This is because in the present model we have not included transport of NTBI into any other organ, and thus all the excess iron stays in the plasma. Together with the experimental results, this suggests that there are likely also other transporters that allow NTBI uptake in other organs.

**Table 2.** ZIP14 knockout on hemochromatosis background. The computational model was modified to suppress hepcidin synthesis (HAMP^-/-^) simulating hemochromatosis, and the rate of liver uptake of NTBI was also set to zero to simulate ZIP14 knockout (HAMP^-/-^:ZIP14^-/-^) and variables were quantified after 365 days. Change is measured as the ratio HAMP^-/-^:ZIP 14^-/-^/HAMP^-/-^.

| Variable | $HAMP^{-/-}$ | $HAMP^{-/-}$<br>$ZIP14^{-/-}$ | Change |
| --- | --- | --- | --- |
| [FeDuo] (mM) | 2.04 | 2.04 | $1.0 \times$ |
| [FeRBC] (mM) | 24.5 | 24.5 | $1.0 \times$ |
| [FeBM] (mM) | 3.35 | 3.36 | $1.0 \times$ |
| [FeSpleen] (mM) | 0.532 | 0.533 | $1.0 \times$ |
| [FeLiver] (mM) | 20.8 | 0.466 | $0.0224 \times$ |
| [NTBI] (μM) | 8.84 | 18,115 | $2049 \times$ |
| [FeRest] (mM) | 0.228 | 0.228 | $1.0 \times$ |
| [hepcidin] (nM) | 0 | 0 |  |
| [EPO] (pM) | 1.60 | 1.60 | $1.0 \times$ |
| TF saturation (%) | 99.8 | 100 | $1.0 \times$ |
| Total Fe (mg) | 2.74 | 2.73 | $1.0 \times$ |

### Anemia of inflammation

Anemia of inflammation is caused by high levels of IL-6 which induces hepcidin secretion by hepatocytes (Nemeth *et al*, 2004; Weiss & Goodnough, 2005) resulting in hypoferremia with increased iron accumulation in tissues, and impaired erythropoiesis (Nemeth *et al*, 2004; Weiss & Goodnough, 2005; Kim *et al*, 2014). Transgenic overexpression of hepcidin in mice causes a phenotype similar to anemia of inflammation (Roy *et al*, 2007), while hepcidin knockout mice (HAMP^-/-^) display a faster recovery from anemia of inflammation (Kim *et al*, 2014), indicating that hepcidin has a central role in this pathology. Using the computational model, we simulated an overexpression of hepcidin to check whether it would result in a phenotype like that of anemia of inflammation. Hepcidin overexpression was simulated by adding one constant term to the rate law for the synthesis of hepcidin in the computational model. This increases the synthesis rate of hepcidin independently of the amount of transferrin-bound iron, effectively increasing a basal synthesis rate for this hormone. The simulation, depicted in Figure 4, resulted in a small reduction of iron in RBC, a significant accumulation of iron in the spleen, lower transferrin saturation and lower [*NTBI*] (hypoferremia), and elevated [*EPO*]. These results are similar to the experimental results of Roy *et al.* (2007). Iron also accumulated in the liver, in line with results from Kim *et al.* (2014). These changes become larger with an increased degree of hepcidin overexpression (Figure S5). The most pronounced effect was the iron accumulation in the spleen. Note that the transient recovery of hepcidin observed is due to the depleted plasma iron as well as to the negative feedback from elevated [*EPO*]. We conclude that the model is able to reproduce the phenotype displayed in anemia of inflammation.

**Figure 4.**
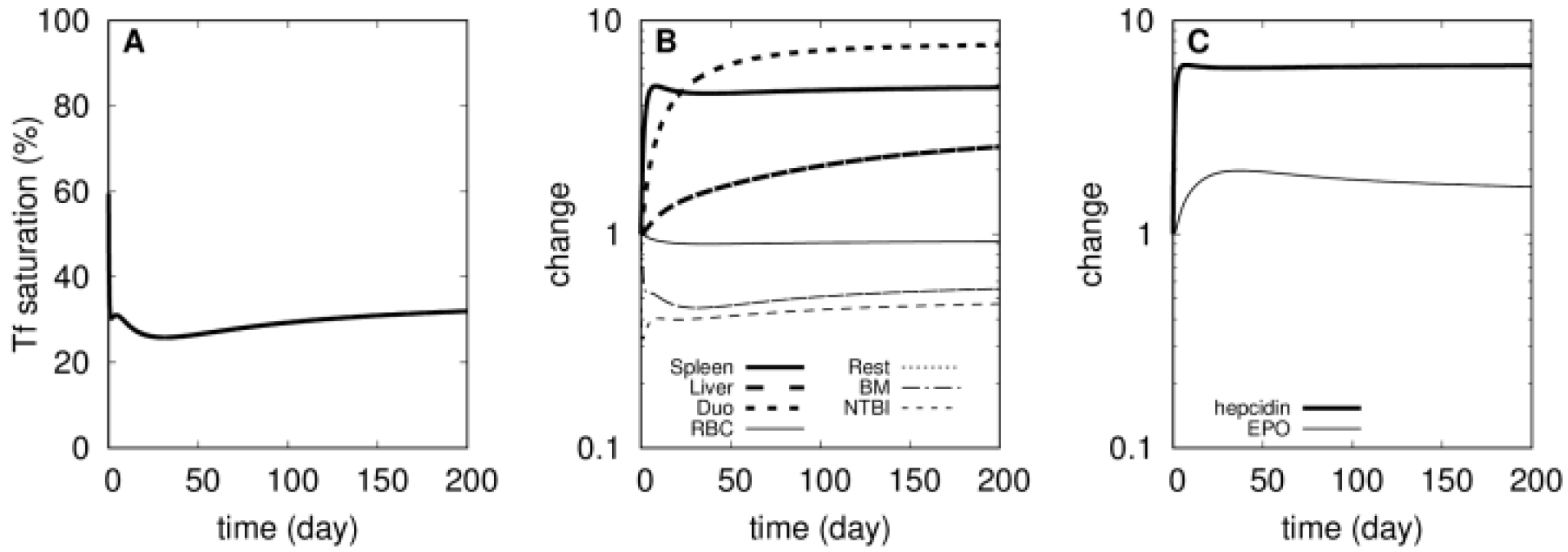
Simulated physiological changes in anemia of inflammation. At time zero the basal rate of hepcidin synthesis is raised (from 0 to 10^−7^ M/day) and the time course is followed for 200 days. A – Transferrin saturation. B – Relative changes in iron content of several compartments (abbreviations: ‘Duo’ for duodenum, BM for bone marrow, ‘Rest’ for rest of body). C – Relative changes in [*EPO*] and [*hepcidin*].

### β-thalassemia

β-thalassemia is a hereditary disease caused by reduced or absent production of hemoglobin’s β-globin chain. This results in a defective erythropoiesis where α-globin tetramers precipitate and cause oxidative membrane damage leading to apoptosis of a large proportion of RBC precursor cells (Origa, 2017). It also results in shorter lifespan of mature RBC due to their increased catabolism by splenic macrophages (Skow *et al*, 1983; Yang *et al*, 1995; Mathias *et al*, 2000). Partial loss of β-globin synthesis results in a milder form of the disease known as thalassemia intermedia while its complete loss results in an extreme form, thalassemia major. The phenotype includes reduced hemoglobin level and RBC count (Gardenghi *et al*, 2007; Ginzburg *et al*, 2009; D’Anna *et al*, 2011), tissue iron overload especially in the liver and spleen (Ginzburg *et al*, 2009; Gardenghi *et al*, 2010), increased plasma iron and transferrin saturation (Browne *et al*, 1997; Ginzburg *et al*, 2009; Gardenghi *et al*, 2010), decreased hepcidin level (Gardenghi *et al*, 2010; Gelderman *et al*, 2015) and increased EPO level (Gelderman *et al*, 2015).

While the present computational model does not represent hemoglobin explicitly, it is still possible to use it to simulate the thalassemia phenotype in terms of the iron distribution. To achieve this the model was modified to have a reduced rate of iron transfer from plasma to the bone marrow (parameter *kInBM*), representing the impaired erythropoiesis. In addition the rate of transfer of iron from RBC to spleen (*kRBCspleen*) was increased, representing the increased destruction of RBC by spleenic macrophages with resulting shorter RBC lifespan. The parameter *kRBCspleen* was increased four fold given the experimental observations of Gelderman *et al.* (2015) that found the RBC half-life four-fold lower in thalassemic mice compared to wild type. By modulating the degree of change in *kInBM* (rate constant for transfer of iron from transferrin to bone marrow) the model could simulate thalassemia intermedia and thalassemia major (Figure 5). This parameter change reflects the efficiency of erythropoiesis in the model, and thus by decreasing it the model is able to reproduce the main features of the thalassemia phenotypes. Figure 5 depicts the changes in phenotype as this parameter is reduced. At around 60% of *kInBM*’s original value there is a qualitative change in the phenotype, suggesting that this may reflect the transition between the two types of thalassemia; this qualitative change seems to depend on the full saturation of transferrin (Fig. 5A).

**Figure 5.**
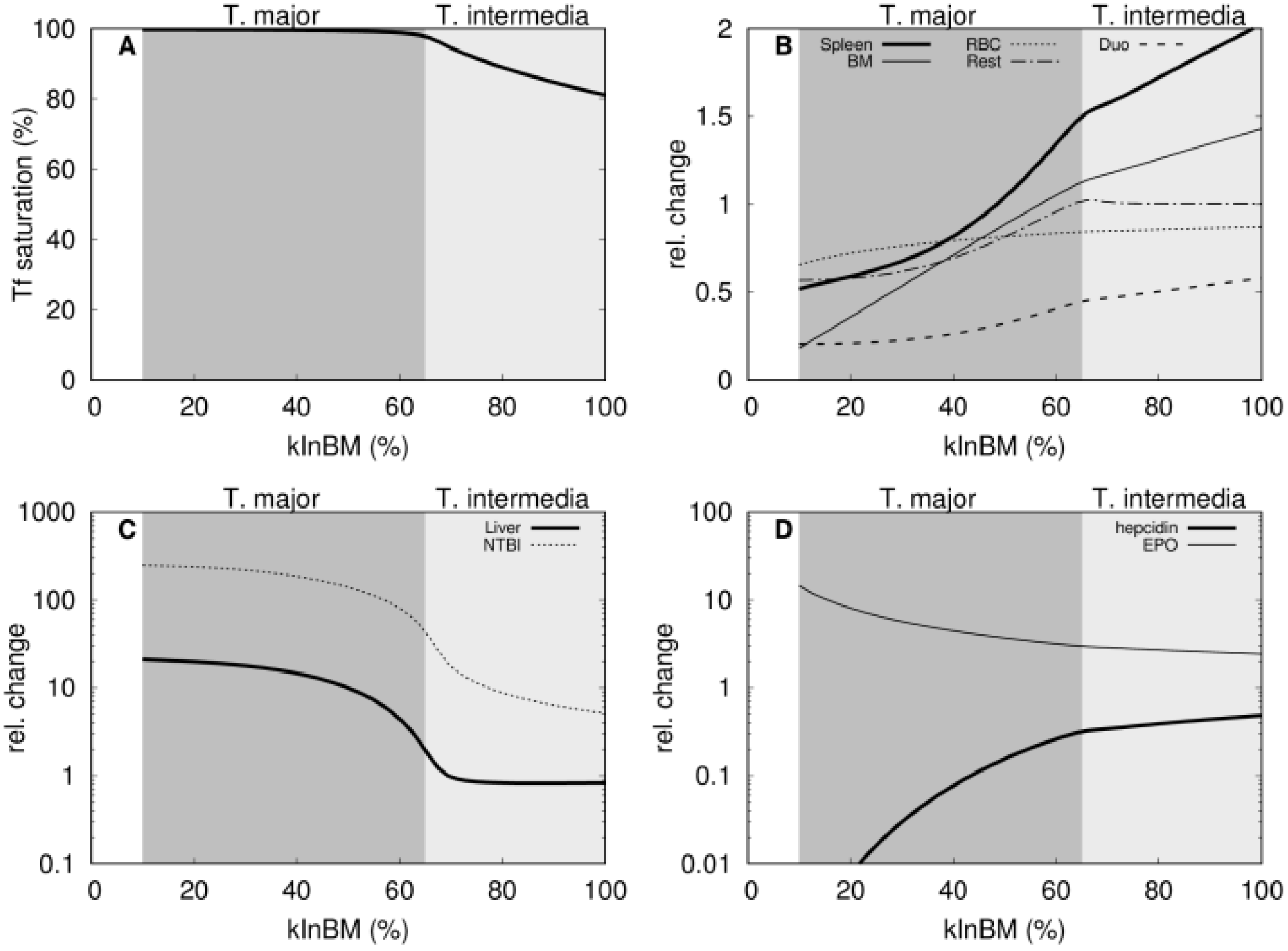
Simulation of thalassemia. The model was changed by increasing the rate of transfer of RBC iron to spleen (*kRBCspleen*) four-fold. The plots then show the effect of slower rates of incorporation of transferrin iron to bone-marrow (*kInBM*). **A**: transferrin saturation; **B**:iron content in spleen, RBC, duodenum, bone marrow, and rest of body; **C**: NTBI and liver iron; **D**: hepcidin and EPO. A qualitative change in the phenotype happens at around 60-65% value of *kInBM*, corresponding to the full saturation of transferrin; for values below this threshold the phentype resembles thalassemia major, whereas above is more like thalassemia intermedia.

For a small decrease of paramater *kInBM, e.g.* 80% of its original value corresponding to mild impaired erythropoiesis, the model shows iron accumulation in the spleen but not in the liver. This pattern of iron accumulation is in good agreement with experimental data pertaining to thalassemia intermedia (th3/+ mice) (Ginzburg *et al*, 2009). The trends in other variables, such as increased NTBI, Tf saturation, and EPO and lower hepcidin are also in good agreement with experimental observations in a thalassemia mouse model (Ginzburg *et al*, 2009; Gardenghi *et al*, 2010; Gelderman *et al*, 2015). By further reducing the parameter *kInBM, e.g*. down to 50% of its original value, the model predicts high iron accumulation in the liver however the spleen iron is back to wild type levels. This is also supported by experimental observations in the thalassemia major mouse model (th3/th3^tp^) at 2 months (Ginzburg *et al*, 2009). Other marked changes are a large increase of NTBI and EPO, and a severe reduction of hepcidin, which are also characteristic of the thalassemia major phenotype (Ginzburg *et al*, 2009).

The model simulation suggests that the most important qualitative change between thalassemia minor and major is the full saturation of transferrin, leading to a complete disregulation of iron homeostasis with extremely high NTBI levels and very low hepcidin. The model is able to simulate these qualitative changes even though the change in the efficiency of erythropoiesis decreases gradually from 80% to 50% of wild type level (Fig. 5A). Note that normally high transferrin saturation would tend to induce hepcidin secretion and this would control the iron level, but the inhibitory effect of EPO on hepcidin (through erythroferrone) overcomes the transferrin-bound iron induction leading to a very low hepcidin level in the thalassemia major phenotype. This in turn increases iron absorption which further increases NTBI and liver iron accumulation.

An important role for computational models is that they allow us to examine the individual contribution of each parameter that may be difficult to elicit independently in experiments. The etiology of β-thalassemia includes two associated parameter changes that are not easily separated in experiments: the degree of erythropoiesis impairment and the extent of RBC lifspan shortnening (both are caused by the excess α-globin in erythroid cells). This, however, is easy to analyze with the model, where each of these perturbations can be applied independently. Thus, Figure 6 shows the independent role of each of these two contributions, as embodied by the *kInBM* and *kRBCSpleen* model parameters. Interestingly it becomes apparent that the RBC lifespan (*kRBCSpleen*) alone is not able to push the phenotype into the thalassemia major domain (as judged by 100% Tf saturation). On the other hand it is always possible to push the phenotype into that range by decreasing the efficiency of erythropoiesis (*kInBM*). This suggests that the inefficient erythropoiesis is a more determinant cause of the disease than the RBC half-life.

**Figure 6.**
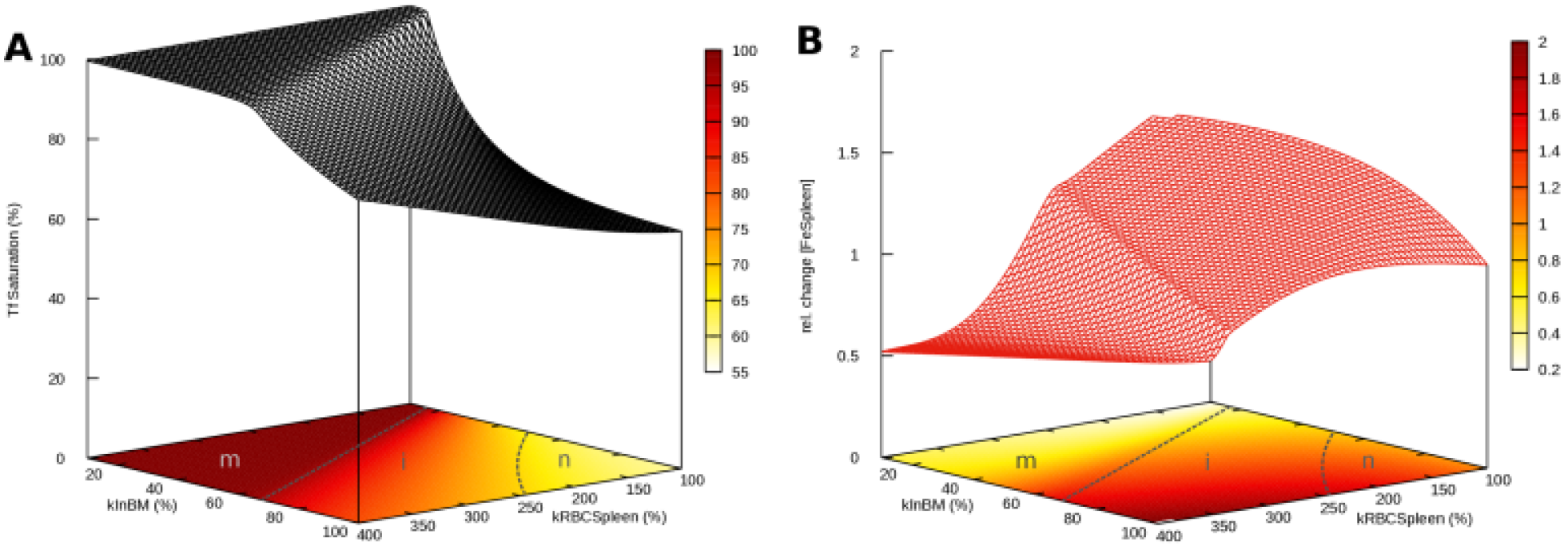
Effect of efficiency of erythropoiesis and RBC lifespan on β-thalassemia phenotypes. Parameters *kInBM* and *kRBCSpleen* were varied and simulated for 365 days. **A**: effect on transferrin saturation; **B**: effect on spleen iron content. The efficiency of erythropoiesis is directly proportional to parameter *kInBM*, while the half-life of RBC is inversely proportional to *kRBCSpleen* (at 400% the half-life is ¼ of its original value). The contour plots at the base include proposed boundaries between thalassemia major (m), thalassemia intermedia (i) and normal (n) phenotypes in dashed lines (the n“i boundary is somewhat arbitrary, while m4 is dictaded by the full saturation of Tf).

### Transferrin treatment of β-thalassemic mice

Transferrin treatment has been shown to reduce ineffective erythropoiesis, improve hemoglobin level, elevate hepcidin expression, and normalize the iron content of liver and spleen in β-thalassemic mice (Li *et al*, 2010; Gelderman *et al*, 2015). We investigated if the computational model could simulate the effect of this treatment accurately. For this purpose, we simulated the experimental protocol where thalessemic and wild type mice were injected with 10 mg of transferrin daily for 60 days (Li *et al*, 2010; Gelderman *et al*, 2015); accordingly the simulation adds 10 mg apotransferrin (equivalent to increasing the Tf plasma concentration by 102 μM) for the same period of 60 days. In the model this periodic Tf treatment applied to the “wild type” model did not affect the iron content of most compartments, which is consistent with the experimental results of Li *et al.* (2010), the only significant changes observed were a decrease in [*NTBI*] (about 10x) with an associated decrease in transferrin saturation (down to 10%). Then we simulated applying the same treatment to a thalassemia major version of the model (*kInBM* at 25% and *kRBCSpleen* at 400% their WT values, see previous section) and this resulted in higher iron level in RBC, lower [*NTBI*], a normalized Tf saturation, normalized liver iron (*i.e*. similar to WT) as well as lower total body iron, compared to the values before treatment. Additionally the concentration of EPO decreased while that of hepcidin increased. All these indicators become closer to their values in the wild type, demonstrating that the model predicts a good response from transferrin treatment as did the experiments (Li *et al*, 2010; Gelderman *et al*, 2015). These results are depicted in Figure 7, where it is also evident that these beneficial effects only last while the transferrin treatment is applied (shaded area); once treatment ceases those variables go back to their diseased levels, suggesting that to be effective this treatment would likely need to be life-long. The only major discrepancy between the simulation and *in vivo* observations is that in the former the spleen iron content increases during the transferrin treatment (Fig. 7F) while *in vivo* this has been observed to decrease (Li *et al*, 2010; Gelderman *et al*, 2015). This increase in the model is likely caused by the increase in hepcidin concentration (compare Fig. 7F and 7G) causing retention of iron in the macrophages. On the other hand, the liver, which is similarly responsive to hepcidin, does not increase its iron level, but rather has it reduced to wild type levels during the treatment (as also seen in the experiments). Analysis of the model strongly suggests that this reduction in the liver is due to to a much lower influx of NTBI through ZIP14 caused by the drastic fall of [*NTBI*] (Fig 7C).

The discrepancy between the simulation and experiments relative to spleen iron is likely due to the fact that *in vivo* the RBC half-life increases with the transferrin treatment (Li *et al*, 2010). This has an effect on the spleen iron level opposite to that of the hepcidin increase, the former likely overcoming the latter. Li *et al.* (2010) attribute the increased RBC half-life to a decrease in α-globin, but since the current model has no representation of hemoglobin it cannot reproduce this effect, other than by changing the value of the *kRBCSpleen* parameter, which directly determines the RBC half-life. Apart from this discrepancy on spleen iron, every other aspect of iron distribution in this proposed thalassemia treatment with transferrin injections is replicated in the model.

Finally the model also shows that while daily injections of Tf are very effective (Fig. 7), lower frequencies of injection may be sufficient to increase the RBC iron level to a similar extent. However, these lower frequencies will lead to oscillatory patterns in the other variables, driven by the half-life of transferrin (Figure S6). When the injections are applied daily, these oscillations are only really detectable in the total amount of transferrin and NTBI (Fig. 7A and B)

**Figure 7.**
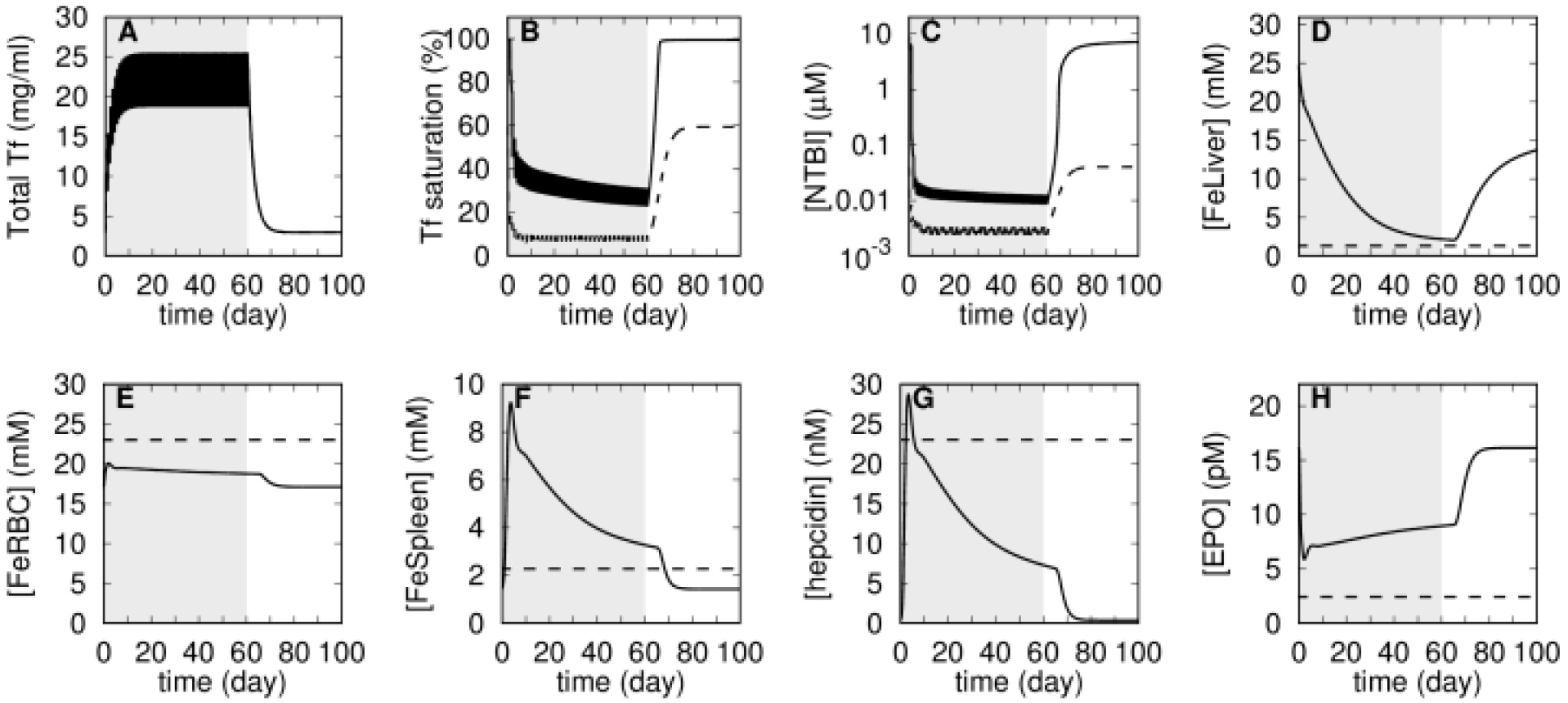
Simulation of transferrin treatment on thalassemia and wild type. Daily injections of apo-transferrin were simulated by increasing apo-transferrin plasma concentration by 102 μM daily for a period of 60 days (shaded area). This procedure is applied both to the wild type model (dashed lines) and to the thalassemia major model (solid lines). The thalassemia major model is initialized by reducing parameter *kInBM* to 25% WT value, and *kRBCSpleen* increased to 400% WT value, then a time course of 365 days is run prior to the application of the transferrin treatment (not shown on this figure). A – total transferrin; B – transferrin saturation; C – NTBI concentration; D – liver iron concentration; E - RBC iron concentration; F – Spleen iron concentration; G – hepcidin concentration; H – EPO concentration.

### Erythroferrone inhibition in β-thalassemia

Kautz *et al.* (2015) showed that an ablation of expression of erythroferrone in thalassemic mice restored the hepcidin level and improved iron overload, however it did not improve the hemoglobin level. In our present model erythroferrone is not represented explicitly but its role is included as an inhibition of hepcidin synthesis by EPO (Equation 6). Simulation of a decreased expression of erythroferrone can be achieved by weakening that inhibition effect (increasing the inhibition constant *KEPOhepcidin*). Thus we compared the model variables when this inhibition has been lowered in the thalassemia simulation (see above). Figure 8 depicts a comparison between the simulation results of the wild type, thalassemia major, and thalassemia major with decreased EPO inhibition of hepcidin. The simulation behaved essentially like the experimental observations, except what concerns the spleen iron level. As in the experiments, decreasing the action of erythroferrone in the simulation causes a normalization of liver iron, NTBI, transferrin saturation and hepcidin levels, and it does not improve the iron content of RBC (or bone marrow). However, contrary to the experiments the model increased the spleen iron content, which is similar to the observations of the previous section, pointing to a part of the model that could be improved. Since the hepcidin level was restored from a low value, this model increase of spleen iron is due to the consequent ferroportin blocking effect; since this effect is not observed *in vivo* it suggests that the spleen iron traffic is likely more complicated than represented in the model (where iron can only enter through transferrin or erythrophagocytosis).

**Figure 8.**
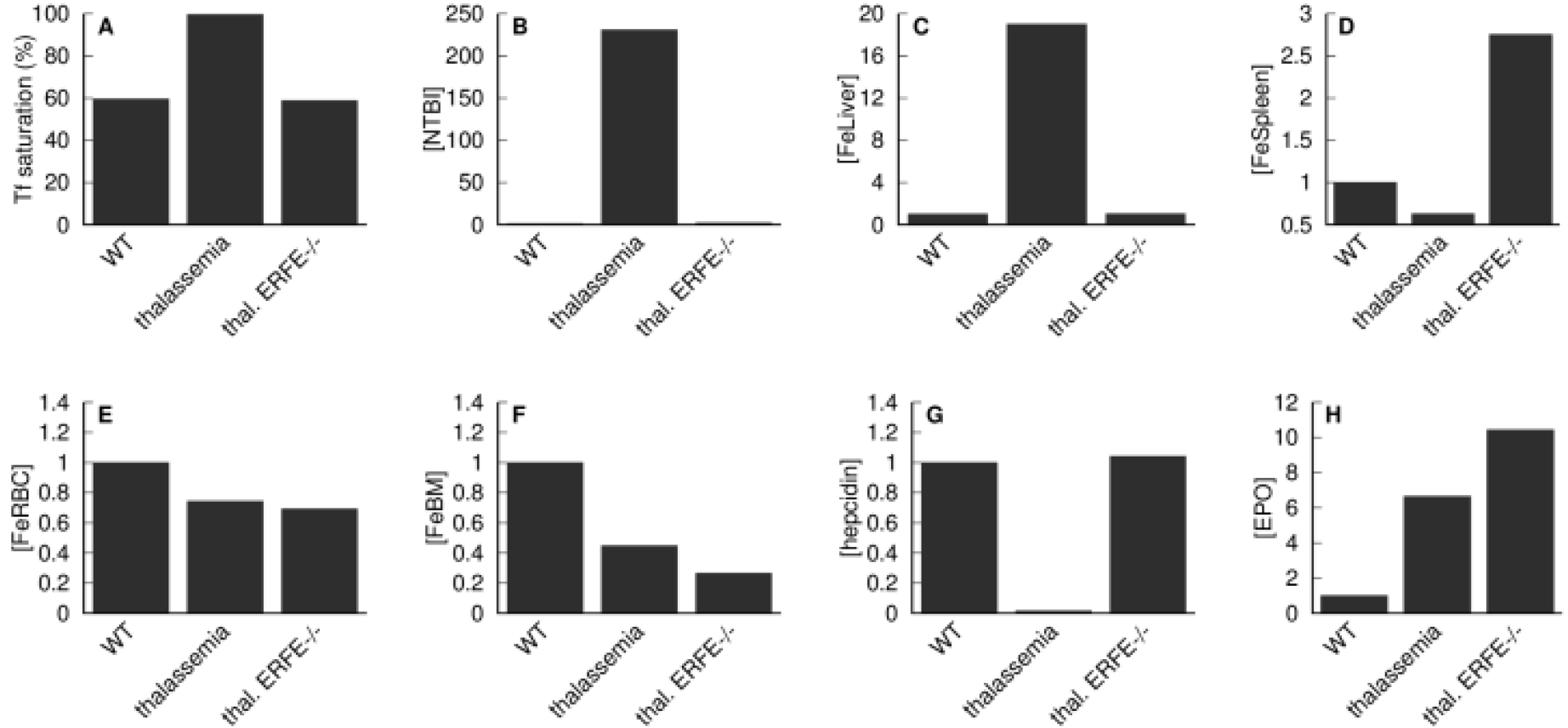
Effect of a reduction of erythroferrone action on thalassemia. The thalassemia major phenotype is compared against the wild type, as well as the phenotype of thalassemia with reduced inhibition of hepcidin synthesis by EPO. The thalassemia major model is initialized by reducing parameter *kInBM*to 25% WT value, and *kRBCSpleen* increased to 400% WT value; the thalassemia with reduced erythroferrone activity (denominated “thal. ERFE-/-”) has additionally an increase by 9 orders of magnitude (×10^9^) of *KEPOhepcidin*, effectively removing the inhibition. Both in the thalassemia and thalassemia with ERFE-/- cases the model is allowed to follow a 365 day time course before measuring the variables. Apart from the transferrin saturation, all values are expressed as a ratio against the wild type value. A– transferrin saturation; B – NTBI; C – liver iron; D – spleen iron; E – RBC iron; F – bone marrow iron; G – hepcidin; H – EPO.

### Hypotransferrinemia

Hypotransferrinemia is a rare iron overload anemia due transferrin deficiency resulting from mutations in the transferrin gene (TF). Key phenotypic changes include substantial iron overload in non-erythropoietic organs, diminished hepcidin levels, and severe iron deficiency anemia (Trenor *et al*, 2000; Ponka, 2002). This phenotype is very similar between humans and the mouse model (Ponka, 2002). As a further validation of the computational model, we simulate hypotransferrinemia by reducing Tf concentration to 2% of the normal level (achieved by a reduction of its rate of synthesis). Figure 9 shows the differences between wild type model and the model of the hypotransferrinemia (Hpx). In agreement with the experimental mouse data (Trenor *et al*, 2000; Bartnikas *et al*, 2011) the Hpx simulation displays reduced iron in the RBC and spleen, decreased hepcidin, and significant accumulation of iron in the liver relative to the WT simulation. Bartnikas *et al.* (2011) proposed that the reduced hepcidin level in Hpx mice was associated with two signals, one from the loss of Tf-mediated activating signal and the other is the inhibitory signal from erythroid regulators. To determine which signal is dominant in suppressing hepcidin, we also created a model where in addition to the reduced Tf we removed the (erythroferrone-mediated) negative feedback of EPO on hepcidin (denominated Hpx ERFE^-/-^). This resulted in a two-fold increase in the hepcidin level, though still far from the WT level but this did not affect the liver iron accumulation nor the other effects of the Hpx mutation. Thus our simulation results indicate that the signal from erythroid regulators (erythroferrone) does not have an influence in the observed phenotype and the major effect is transmitted through abrogated Tf-mediated signaling.

We also wanted to follow the dynamics of plasma iron in the Hpx mutation. This was achieved through a simulation of an injection of a bolus of ^59^Fe tracer, following its distribution through 365 days. Figure 10 displays the simulations obtained with the Hpx and wild type models. In Hpx the majority of the tracer goes to the liver and is retained there for the entire period of time, while in the wild type only a small proportion of tracer appears in the liver and rapidly decreases. In the wild type a large proportion of the tracer quickly enters the RBC (circa 60%), while in the Hpx it only reaches 20%. Overall our model is able to qualitatively simulate the Hpx phenotype and displays an appropriate organ distribution of radioactive iron under this condition. This provides further validation of the present model.

**Figure 9.**
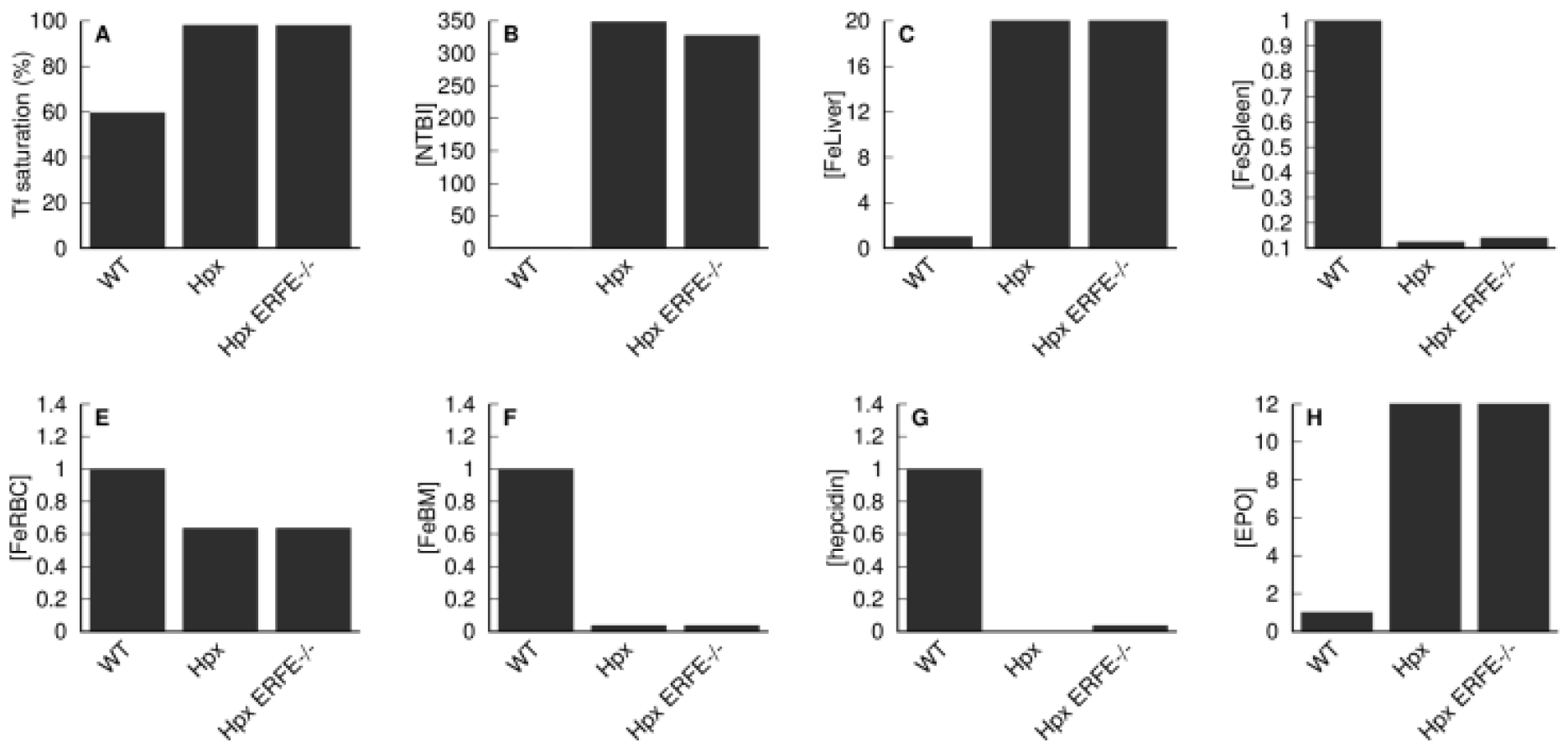
Effect of the reduction of total transferrin (Hpx mutation) on iron levels. Iron and hormone levels are compared between the wild type model (WT) and a model with 2% of WT total transferrin concentration (Hpx), and a model that also has reduced inhibition of hepcidin synthesis by EPO (Hpx ERFE^-/-^). The Hpx model has the *V*_*max*_ for production of transferrin (parameter named *vTF*) at 0.31 μmol/day versus 155 μmol/day in the WT; the Hpx ERFE^-/-^ model additionally has a reduced erythroferrone activity achieved by increasing the parameter *KEPOhepcidin* by 9 orders of magnitude, effectively removing the inhibition of EPO on hepcidin synthesis. Both in the Hpx and Hpx ERFE^-/-^ cases the model is allowed to follow a 365 day time course before measuring the variables. Apart from the transferrin saturation, all values are expressed as a ratio against the wild type value. A– transferrin saturation; B – NTBI; C – liver iron; D – spleen iron; E – RBC iron; F – bone marrow iron; G – hepcidin; H – EPO.

**Figure 10.**
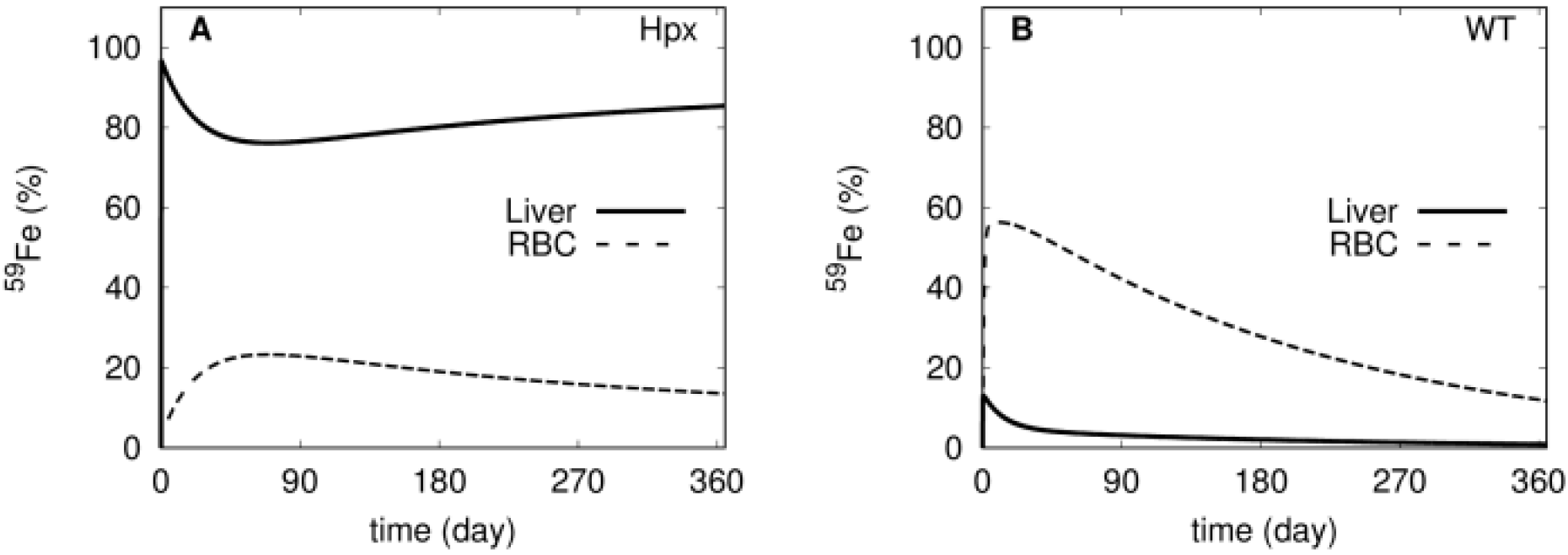
Simulation of ^59^Fe tracer distribution in Hpx and WT models. The amount of tracer is followed for 365 days after injection of ^59^Fe in the plasma (as NTBI* species). A – Hpx model, B – wild type model.

### Comparison of intravenous and intragastric iron administration

Kim *et al. (2013)* investigated the differences between uptake and distribution of iron when administered through intragastric (IG) and intravenous (IV) routes. They observed that when ^59^Fe was administered by IG, the amount of iron accumulated in the blood of hemochromatosis (HFE^-/-^) mice was significantly higher compared to that of wild type mice, while the opposite was observed when ^59^Fe was given by IV route. The present model was also used to simulate these experiments successfully. In this case the simulation of hemochromatosis is achieved by silencing hepcidin synthesis at the transcription level (HAMP^-/-^) instead of the HFE^-/-^ mutation. The model simulations behave in the same way as the experimental results, also showing that IG administration results in higher blood iron accumulation in HAMP^-/-^ than in WT; while the opposite pattern is observed for the IV route (see Figure 11A-B and compare with Figure 2 in Kim *et al.* (2013)). The measurements reported by Kim *et al.* (2013) were for total blood radioactivity, which includes NTBI, transferrin-bound iron, and RBC iron, and this does not easily allow for a clear explanation of the phenomenon, but those authors suggested that it could be due to an increased clearance of NTBI from the plasma in hemochromatosis, compared with the wild type (Kim *et al*, 2013). This is where a model can be uniquely useful to aid in the interpretation of results since in the model all variables are readily available and it is possible to carefully inspect the fate of the tracer. Figure 11C-D depicts the simulated dynamics of the administered ^59^Fe in the first four hours in the three components of blood (NTBI, Tf, and RBC), as well as the liver compartment. It is clear that the major difference is that in the HAMP^-/-^ the liver is rapidly accumulating a large proportion of the tracer. This explains the differences observed in the case of IV administration: in HAMP^-/-^ the tracer is moving from NTBI to the liver at a fast rate, leaving less tracer in the blood than in the WT. But this is not the reason why there is more blood iron in the HAMP^-/-^ when the tracer is administered through IG. To explain this we have to look at the duodenal rate of absorption of the tracer (*i.e*. the flux of tracer from the duodenum into the plasma). Fortunately the model also readily provides the fluxes, depicted in figure S7, and from those data it becomes clear that HAMP^-/-^ absorbs the tracer at a much higher rate than the wild type, and that flux is higher than the flux of tracer binding to transferrin. This stems from the absence of hepcidin, which results in a higher activity of duodenal ferroportin and consequentely higher iron absorption.

While the simulation differs slightly from the experiment, by causing hemochromatosis through mutation in the HAMP gene (hemochromatosis type 2B) rather than through the HFE gene (hemochromatosis type 1), the results are qualitatively similar to the experiments. The major difference between the two is the complete absence of hepcidin in the model versus a low level of hepcidin level in the experiment, though clearly both result in a similar phenotype. This example illustrates the benefits of using computational modeling and simulation with a validated model to aid the interpretation of experimental results.

**Figure 11.**
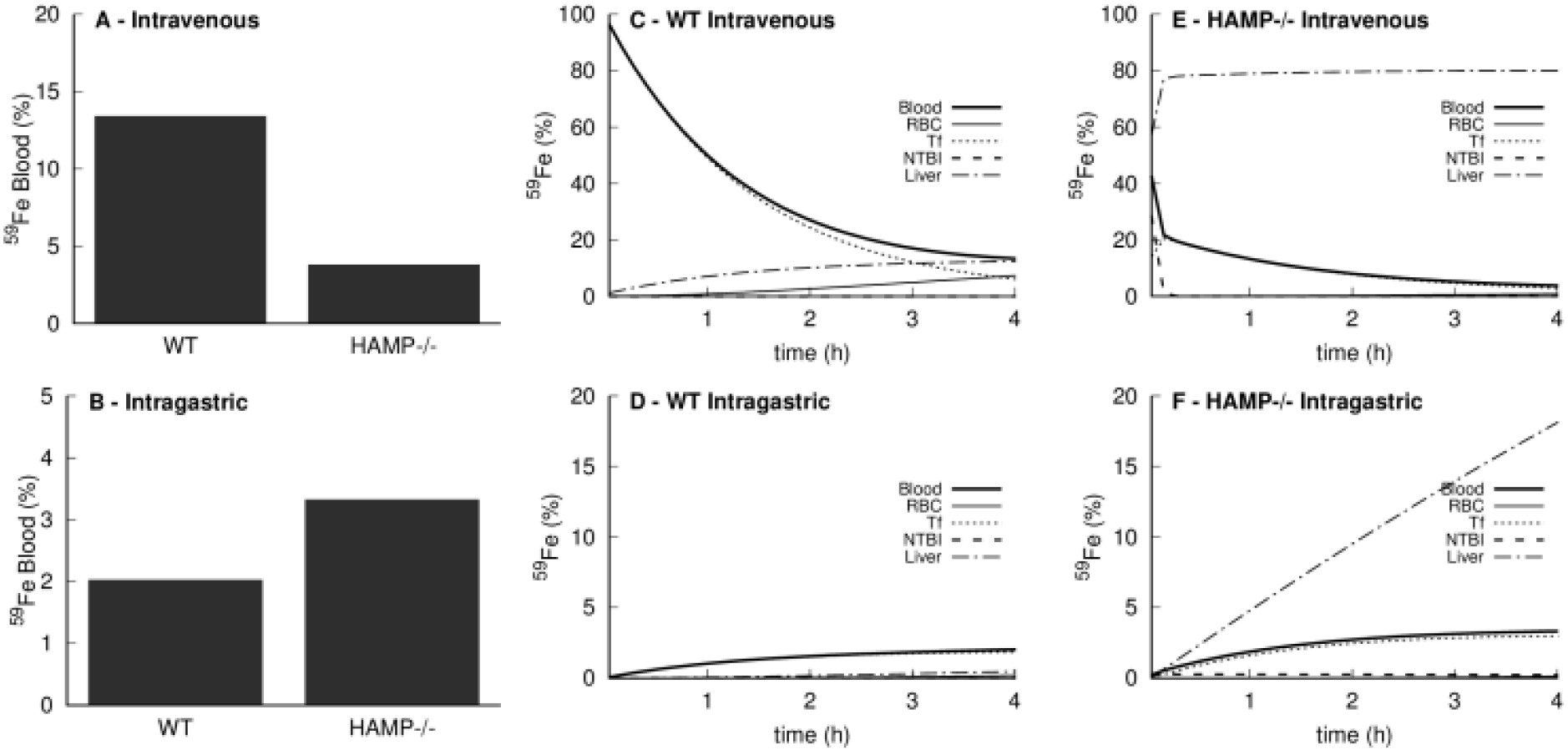
Simulation of iron tracer administration by intravenous and intragastric routes. A bolus of iron tracer is administered in the plasma as radioactive NTBI (simulating an intravenous injection) or as radioactive duodenal iron (simulating intragastric gavage). Total blood iron tracer is calculated as the sum of NTBI tracer, trasnferrin bound tracer, and RBC tracer. **A-** percentage of tracer present in blood four hours after intravenous injection in the wild type and HAMP^-/-^; **B-**percentage of tracer in blood four hours after intragastric gavage in wild type and HAMP^-/-^; **C-**tracer time course in the wild type intravenous injection; **D-** tracer time course in the HAMP^-/-^ intravenous injection; **E-** tracer time course in the wild type intragastric gavage; **F-** tracer time course in the wild HAMP^-/-^ intragastric gavage.

## Discussion

A mathematical model that integrates our current knowledge on iron metabolism would provide valuable information to better understand its homeostatic regulation, and it could be an aid to experimental design and interpretation of results. Previously, we developed such a model centered on hepcidin regulation that successfully reproduced iron dynamics under iron“adequate and iron“deficient diets but not the dynamics under an iron“rich diet (Parmar *et al*, 2017), failing to reproduce iron accumulation in the liver and overestimating RBC iron under iron“rich conditions. While that model could simulate anemia of chronic disease, it was unable to reproduce the phenotype of hereditary hemochromatosis appropriately. The results from that earlier model indicated that hepcidin regulation alone was not sufficient to explain phenotypes under high iron conditions, and suggested that other mechanisms must play a significant role. Another recent modeling study on iron regulation under inflammation (Enculescu *et al*, 2017) also fails to explain high iron conditions and overestimates RBC iron in the simulation of hemochromatosis, unlike well“established experimental results that show RBC iron to be essentially unaltered in that condition (Zhou *et al*, 1998).

Here, we presented a new mathematical model that can accurately explain the iron distribution under low and high iron conditions. The model required the addition of two important mechanisms: 1) NTBI uptake by the liver, and 2) the regulation of erythropoiesis by erythropoietin. First, we calibrated the model using the same ferrokinetic time“series data (Schümann *et al*, 2007; Lopes *et al*, 2010) used to calibrate previous models (Parmar *et al*, 2017; Enculescu *et al*, 2017). Then the model was validated by simulating several mouse experiments related to pathological conditions that affect iron dynamics and regulation. This included hemochromatosis, β-thalassemia, anemia of inflammation, and hypotransferrinemia. Importantly these simulations did not attempt to fit parameter values to the data, they simply replicated an independent set of experiments still using the same parameter set. This extensive validation provides a strong confidence in the predictive powers of the model.

One aspect where computational models can be very helpful is in asking certain *what if?* type questions that are impossible to execute experimentally. Here we used the model to investigate the relative contribution of the two major mechanisms causing the β-thalassemia pathophysiology: inefficient erythropoiesis and lower RBC half-life. Experimentally these are both consequences of an unbalanced composition of hemoglobin types and are not easily separated. In the model they can be manipulated independently and doing so suggested that the inefficiency of erythropoiesis is the largest contributor to the pathophysiology. Lower RBC half-life alone would never lead to the observations typical of thalassemia major, but impaired erythropoiesis could. We believe that this use of the model to better understand the underlying causative biology is perhaps its most important role, functioning as a kind of “executable” and quantitative review of the underlying biology.

The application of the model to simulate transferrin treatment in thalassemia provides an example of how such a model could eventually be used to plan and optimize therapies and interventions. The model was used to investigate the effect of the frequency of Tf injections from daily to every four days; the results in Figure S6 show that injections every other day would have almost the same effect as daily injections. In terms of just decreasing the liver iron content, even injections every four days would have a similar efficiency. The present model can then be useful to help design certain mouse experiments. Obviously, for therapeutic use the model would have to be adapted and validated with human data. If such a human model could also be calibrated for specific subjects it would then become a useful tool for personalized medicine.

The model replicated the experimental observation that when Tf saturation is high the organ distribution of injected iron is different than when it is lower (Figure 3). Future human personalized models would allow one to design better interventions for anemic patients where the amount of injected iron would not exceed that person’s transferrin capacity (calculated from the model), maybe allowing for an optimized multiple injection schedule. This would maximize the transfer of the injected iron to the erythropoietic system instead of its accumulation in the liver or other organs causing side effects.

The strategy behind the development of this model is that it ought be as simple as possible to explain the iron distribution in the whole body focusing on the most relevant compartments. This meant selecting the important compartments: liver, bone marrow, spleen, duodenum and all components of blood, namely red blood cells, transferrin-bound iron and non-transferrin bound iron. The rest of the body is still accounted for in quantitative terms, but aggregated in a single compartment. Further models could benefit from increased compartmental resolution (e.g. separating the brain, pancreas or kidneys from the “rest of body” compartment). Another important simplifying criterion was to limit intracellular regulation mechanisms to a minimum. Thus the iron regulatory protein-iron regulatory elements system (IRP-IRE, Anderson *et al*, 2012) was not included here, nor was there a distinction made between different intracellular iron states (labile iron pool, ferritin-bound, enzyme-bound, etc.) Interestingly, the model is still able to largely follow iron distribution dynamics at the organ level, thus showing that those intracellular molecular details may not be so important at the whole body level – a finding that is perhaps surprising. But some molecular details did have to be included: a saturable ferroportin-mediated export of iron and its inhibition by hepcidin (Ganz, 2005), and liver import of NTBI through Zip14 (Liuzzi *et al*, 2006; Jenkitkasemwong *et al*, 2015) revealed to be crucial molecular details that impact whole-body iron distribution. The case of Zip14 is well illustrated in the result sections on hemochromatosis, thalassemia, and the differential dynamics of iron upon intravenous injection or intragastric gavage. We had previously ignored the effect of erythropoietin regulation (Parmar *et al*, 2017), but this erythropoiesis regulatory system also proved to be essential to explain the physiology of high iron conditions. While the model does not explicitly consider the recently discovered hormone erythroferrone (Kautz *et al*, 2014), its action is included as an inhibition term between the level of EPO and the synthesis of hepcidin (Equation 6).

The extensive validation passed by this model gives a very good level of confidence in its predictive powers, but it is also important to discuss its limitations. When applying the model to conditions where NTBI import to the liver becomes significant, we noted that the model would not reach a steady state, instead accumulating unlimited amounts of iron in the liver. A steady state would likely be reached if the model included ferritin and its regulation by the IRP system. On the other hand, it is unknown how much iron can be accumulated in the liver. It so happens that the most common complication from untreated hemochromatosis is liver disease caused by severe iron accumulation, so perhaps this lack of a steady state is not so problematic and does reflect the pathology. The model only considers direct import of NTBI to the liver, however there is evidence that this happens also for other organs like the heart and pancreas (Craven *et al*, 1987; Jenkitkasemwong *et al*, 2015). Finally the model does not fit the spleen data very well (see Figs. S3-S4) and some of the validation were also less successful for the spleen than for the other organs. This is likely due to the fact that the model does not consider the erythropoietic function of the spleen, nor does it consider that, in addition to spleen macrophages, the RBC are also degraded by liver Kupffer cells. To overcome this would require adding more detail in these two organs, leading to a multi-scale model that considers different cell types in each organ. Note, however, that this deficiency is quantitatively small.

Given the strong emphasis on the ability of this model to reproduce high iron conditions, it is perhaps opportune to speculate about the role of the liver in taking up iron. While developing the model to support high iron conditions it became obvious to us the direct import of NTBI into the liver was required in high plasma iron concentrations but should not operate under lower plasma iron concentrations. This means that the liver only takes in NTBI when it is in excess in the plasma, and this has been confirmed experimentally (Jenkitkasemwong *et al*, 2015) and replicated by the model (Fig. 3). An interesting question is what mechanism leads to an increased activity of Zip14 in high iron conditions? In the model this was simplified by adopting a kinetic rate law for Zip14 with substrate activation, thus its activity only becomes significant at high substrate concentration ([*NTBI*]). But in reality Zip14 is known to be induced by iron overload (Nam *et al*, 2013), but it is not clear how. The mechanism must depend either on NTBI or transferrin-bound iron because the Hpx mice lacking transferrin accumulate large amounts of iron in the liver (Trenor *et al*, 2000; Bartnikas *et al*, 2011), supposedly due to high expression level of Zip14. So what is the pathway for Zip14 induction under iron overload? What is the signal? What is the receptor? These are interesting questions that ultimately need be addressed with new experiments but the present model could also contribute by allowing to test hypotheses prior to experimentation.

In summary we presented a whole-body model of iron dynamics for the mouse that is consistent with a wide range of dietary iron intake. The model was subsequently validated with a series of independent experiments showing that it behaves very closely to all their results. We demonstrate how the model can be applied to better elucidate the operation of mechanisms, applying it to the comparative contribution of erythropoiesis and RBC half-life to thalassemia symptoms. The model was also used to show how it could have a role in planning interventions (transferrin treatment in thalassemia). We discussed aspects of the model that can still be improved and speculate on how a human specific model could have a role in personalized medicine.

## Acknowledgements

We thank Tom Bartnikas, Karin Finberg, Tim Barry, Mary J. Gockenbach, and Arlie Koziol for discussions about the model. PM thanks the National Institute for General Medical Sciences for funding of COPASI development (GM-080219).

